# Bio-based decontamination and detoxification of total petroleum hydrocarbon contaminated dredged sediments: perspectives to produce constructed technosols in the frame of the circular economy

**DOI:** 10.1101/2023.06.18.545458

**Authors:** Simone Becarelli, Giacomo Bernabei, Giovanna Siracusa, Diego Baderna, Monica Ruffini Castiglione, Giampiero De Simone, Simona Di Gregorio

**Affiliations:** Department of Biology, University of Pisa, Italy; BD Biodigressioni srl, Pisa, Italy; Chemservice S.r.l. - Lab Analysis Group, Via F.lli Beltrami, 15, 20026, Novate Milanese, Milano, Italy

**Keywords:** mycoremediation, *Lambertella* sp. MUT 5852, predictive functional metagenomic analysis, *Kocuria* sp., *Sphingobacterium* sp..

## Abstract

To accelerate the depletion of total petroleum hydrocarbons, a hydrocarburoclastic ascomycetes, *Lambertella* sp. MUT 5852, was bioaugmented to dredged sediments co-composting with a lignocellulosic matrix. After only 28 days of incubation, a complete depletion of the contamination was observed. The 16S rDNA metabarcoding of the bacterial community and a predictive functional metagenomic analysis was adopted to evaluate potential bacterial degrading and detoxifying functions. A combination of toxicological assays on two eukaryotic models, the root tips of *Vicia faba* and the human intestinal epithelial Caco-2 cells, was adopted to assess the robustness of the process not only for the decontamination but also for the detoxification of the dredged sediments. Bacterial taxa, such as *Kocuria* and *Sphingobacterium* sps. resulted to be involved in both the decontamination and detoxification of the co-composting dredged sediments by potential activation of diverse oxidative processes. At the same time, the *Kocuria* sp. showed plant growth promoting activity by the potential expression of the 1-aminocyclopropane-1-carboxylate deaminase activity, providing functional traits of interest for a technosol in terms of sustaining primary producer growth and development.

## 1. Introduction

Nowadays more than ever, saving soil, a non-renewable natural resource, is mandatory. Among others, the opportunity to decrease its consumption by the production of substrates with comparable properties, deriving from processes of transformation of renewable resources, is promising. In this direction, strategies to produce constructed technosols offer interesting possibilities. Constructed technosols consist in man-made soils containing large amounts of organic and inorganic not toxic wastes or recycled material, such as sewage sludge, compost, or residues from construction (IUSS Working Group WRB 2015). The ongoing increasing urbanization determines an exponential increase in natural soil exploitation by grabbing and degradation (FAO 2015). In this context, EU politics are promoting and financing the recovery of brownfields by the design of sustainable approaches for the rescue of damaged soil matrices. *In-situ* technologies for soil recovery are emerging but they are difficult to control and implement. On the other hand, *ex-situ* treatments and the consequent soil excavation and replacement with clean one from uncontaminated area, are currently adopted. Constructed technosols might substitute the above-mentioned clean soils and encounter the interest of the technical and scientific community, committed to select and promote the sustainability of the interventions for the recovery of brownfields. In the concept of constructed technosol production, is consolidated the one of the recovery of wastes. In accordance with the circular economy, ones their toxicological safety is ascertained, because of chemical physical characteristics or decontamination processes, wastes are classified as recycled materials and might find exploitation to produce constructed technosols (Moraga et al. 2019). The main criteria for the selection of not harmful wastes as recycled materials are i) the not limited availability, ii) absence of toxicity, iii) agronomic properties if exploited to produce constructed technosols (Cascone et al. 2019).

In relation to quantity and availability of wastes to be sustainably recovered, dredged sediments constitute a case of interest. Waterway logistic generates costs for the society because of dredging activities, producing massive amounts of sediments, mainly contaminated by total petroleum hydrocarbons (TPH), deriving from spillage of shipyard’s fuel. In Europe, between 100 and 200 million cubic meters of waterway sediments are dredged per year. Since the 1st of June 2012, because of the implementation of the Waste Framework Directive 2008/98/EG, the reuse of dredged sediments was subject to the end-of-waste criteria and sustainable technologies for their recovery and/or decontamination became mandatory. Bio-based processes have been proposed (Dell’Anno 2020). Among others, the mycorememediation approach, based on the bioaugmentation of hydrocarburoclastic fungi to TPH contaminated sediments co-composting with lignocellulosic matrices, ends up with a matrix that might offer the characteristics to be exploited in the production of technosols (Becarelli et al. 2019, 2021; Di Gregorio et al. 2016). However, a toxicological evaluation of the final product of the bio-based process is still undervalued. The assumption might constitute a critical development for the technology, since, especially in the case of organic contamination, it is difficult to define the trend of degradation pathways of different class of pollutants that, microbially transformed, generate not known intermediates of degradation, difficult if not impossible to be identified and qualified in relation to toxicity. Noteworthy, these intermediates have been reported as even more toxic than the parental pollutants (Riser-Roberts 1998) and might persist at the end of a bioremediation process (Ruffini Castiglione et al. 2016). The sole chemical characterization of known pollutants might be not sufficient to evaluate the toxicity of environmental matrices and to guaranty the sustainability of even a bio-based intervention. In fact, sustainability is a complex concept that is related not only to costs but also to avoidance of negative pressure on the natural environment. On the other hand, toxicological assays are strong instruments to provide data on the bioavailability and toxicity of both the residual primary pollutants and their intermediates of degradation. The ISO 292000:2013 (ISO 29200:2013) is a consolidated assay for evaluating the genotoxicity of soils or soil materials on secondary root of *Vicia faba*. In particular, it is designed to measure and quantify chromosome breakage due to dysfunction of the mitotic spindle and the formation of micronuclei in cells of root tips. The assay is designed to evaluate the genotoxicity not only of soils but also of compost and fertilizers, being of interest to evaluate the possible genotoxicity of recovered matrices to produce technosols. The combination of assays on different eukaryotic models might increase the sensibility of the monitoring. In this context, the adoption of assay on the in vitro model Caco-2 cells is of interest. Caco-2 cells are used as an *in vitro* intestinal epithelial model to evaluate the bioavailability of trace elements from different matrices (Balimane et al. 2000), and the cytotoxicity assay with these cells might be important to explore the bio accessibility and bioavailability of eventual residual toxicity of treated matrices, especially if they are subjected to produce elutriates, endangering water bodies for human consumption.

The scope of this experimentation was the optimization of a co-composting process designed to decontaminate dredged sediments from total petroleum hydrocarbons (TPH, 2378 ± 79 mg/kg on a dry weight base ratio). The sediments were pretreated by soil washing to remove salinity and inorganic contamination. The optimization of the co-composting process was related to the result obtained in a previous experimentation (Becarelli et al. 2021) and was consistent with the decrease of the fungal biomass to be bioaugmented (50% of reduction on a weight base ratio), to increase the sustainability of the approach in terms of costs. The study aimed to evaluate the kinetics of TPH degradation and the response of the bacterial community to the treatment, to envisage co-metabolic transformation capacities of the latter and its involvement in TPH degradation in collaboration with the fungal strain, whose metabolism was monitored by the quantification of ergosterol as marker of the active fungal metabolisms (Buiarelli et al. 2013; Olsson et al. 2003). To the scope a predictive functional metagenomic approach was adopted. The same approach was adopted to evaluate other functional traits of interest for the bacterial community of the co-composting sediments. In particular, the capacity of the bacterial community to sustain plants in coping with environmental stress, mediated by the 1-aminocyclopropane-1-carboxylate (ACC) deaminase activity, involved in the inhibition of the synthesis of the plant hormone of stress, the ethylene (Penrose et al. 2003). This bacterial functional trait is of interest in technosols that are supposed to offer a matrix capable to sustain the growth of primary producers. The potential increment in enzymatic activities involved in the soil carbon cycle was also evaluated by evaluating the bacterial contribution to the dye-decolorizing peroxidase activity (DyP), capable of extracellular oxidation of phenolic and non-phenolic lignin model compounds (Chen et al 2016; Chen et al. 2015). Bacterial strains capable expressing the DyP have been described as potentially involved in the mobilization of not easily bioavailable carbon sources in soil, participating to the transformation of the soil organic matter and consequently to the soil carbon cycle (Bearelli et al. 2021b). In addition, a combination of toxicological assays on eukaryotic models such as the root tip of *Vicia faba* and the human intestinal epithelial Caco-2 cells was adopted to assess the detoxification of the decontaminated dredged sediments, for an eventual their safe exploitation in other sectors, such as the one of the productions of technosols.

## 2. Materials and Methods

### 2.1 TPH Contaminated Dredged Co-composting sediment: Co-composting sediment, Fungal Strain and Chemicals

The contaminated co-composting sediments were dredged in the Navicelli Channel, Pisa, Italy (4341055.90” N; 1022050.80” E). One m^3^ of dredged co-composting sediment were treated with a pilot sediment-washing plant, using tap water as the washing carrier. The procedure was designed to decrease the content in heavy metal and to desalinate the brackish sediments. The characteristics of the washed dredged sediment are reported in Table 1. The fungal strain, *Lambertella* sp. MUT 5852, had been previously isolated from the same site [6]. All the chemicals used were of analytical grade and purchased from Merck (Milan, Italy).

**Table 1.** Chemical/physical characteristics of the dredged sediment

| Components | Dredged sediments |
| --- | --- |
| Total petroleum hydrocarbons | 2378 ± 79 mg/Kg |
| Total phosphate | 15 ± 0.2 mg/Kg |
| Total nitrogen | 0.31 ± 0.02 mg/Kg |
| Chloride | 2.4 ± 0.02 g/L |
| Pb | 21 ± 0.1 mg/Kg |
| Cd | 15 ± 0.2 mg/Kg |
| Zn | 7 ± 0.1mg/Kg |
| Cr | 9 ± 0.2 mg/Kg |

### 2.2 Mesocosms-Scale Experimentation

A total of 18 experimental replicates (mesocosm glass pots), each holding 2 kg of dredged sediments, were prepared, and kept in a temperature-controlled (17 ± 1 ◦C) dark chamber at 60% soil maximum water-holding capacity (WHCmax = 9.6% dry mass). The eighteen mesocosms were co-composted with 20% on a weight-based ratio of lignocellulosic ma-trices (wood chip). A total of 9 of these latter were bioaugmented with fresh biomass of *Lambertella* sp. MUT 5852 grown in ME medium (malt broth, 20 g; yeast extract, 5 g in 1 L H_2_O) at 5% on a fresh-weight-based ratio; 9 mesocosms were inoculated with the same biomass after autoclavation (121 ± 1 °C, 1 Atm, for 40 min). All pots were routinely manually mixed every 3 days of incubation and checked for water content by weighting the pots to keep constant values. A total of three mesocosms amended with *Lambertella* sp. MUT 5852 and 3 mesocosms amended with the autoclaved fungal biomass were sacrificed and analyzed for TPH content at the time of mesocosm assemblage (T0), after 18 (T18) and 28 (T28) days of incubation. Three mesocosms for experimental condition showing TPH depletion were measured for ergosterol, humic and fulvic acid content.

### 2.3 Quantification of TPH, Humic and Fulvic Acids, Ergosterol

Quantification of TPH in the co-composting sediment, quantification of humic and fulvic acids and ergosterol was performed as described in (Becarelli et al 2021a). The D’Agostino and Pearson omnibus normality test was adopted for the quantification of the residuals Quantification of Total Petroleum Hydrocarbons, Humic and Fulvic Acids, and Ergosterol.

### 2.4 Metabarcoding Analysis

The total DNA from the co-composting sediments was extracted using a FastPrep 24™ homogenizer and FAST DNA spin kit for soil (MP Biomedicals), starting from 500 mg of sample, according to the manufacturer’s protocol. The quantity of DNA was measured using a Qubit 3.0 fluorometer (ThermoFisher Scientific, Milan, Italy). The DNA purity and quality was measured spectrophotometrically (Biotek Powerwave Xs Microplate spec-trophotometer, Milan, Italy) by measuring absorbance at 260/280 and 260/230 nm. A total of 200 ng of DNA was used to produce paired-end libraries and for sequencing the V4–V5 hypervariable regions of the bacterial 16S rRNA gene by using the 515F forward primer (5’-GTGCCAGCMGCCG CGGTAA-3’) and 907R reverse primer (5’-CCGTCAATTCCTTTGAGTTT-3’) as primers. The libraries for Illumina sequencing were prepared by Novogene using NEBNext Ultra DNA Library Prep Kit, following the manufacturer’s recommendations, and index codes were added. The library was se-quenced on an Illumina platform by Novogene (25 Cambridge Science Park, Milton Road, Cambridge, CB4 0FW, United Kingdom), and 250 bp paired-end reads were generated.

### 2.5 Data Analysis

Cutadapt plugin for Qiime2 was used to demultiplex and trimm the paired-end reads. Qiime2 v 2022.2 standard pipeline was used to assemble forward and reverse reads, filter their quality and chimeras and to assign them to amplicon sequence variants (ASVs). ASVs clustering was performed using DADA2 workflow implemented in Qiime2, with the classifier trained on the V4–V5 hypervariable region extracted from the Silva 138 99% 16S sequences database. To allow comparison between different samples, ASV abundance per sample data were normalized by rarefaction to the same coverage (99.5% of observed species). Next analyses of the canonical correspondence analysis (CCA), and related statistical tests were performed on R 4.1.2 using Phyloseq, Vegan, and Pheatmap packages (versions 1.37.0, 2.5-7 and 1.0.12), respectively. Non-parametric statistics (Kruskal–Wallis test and related post-hoc tests) on chemical data were performed by ggpubr (version 0.4.0.999). The functional metagenomic prediction for the bacterial community was inferred using PICRUSt2 v. 2.4.1 for unstratified and stratified metagenome contribution based on EC numbers. The KEGG pathway and EC contributions were filtered from the output data of PICRUSt2 v. 2.4.1 and processed by R 4.1.2 (Douglas et al. 2020). The analysis of compositions of microbiomes with bias correction (ANCOMBC) (Lin et al 2020) was performed on both tax-onomical and predicted functional data by R 4.1.2 using ANCOMBC v. 1.6.2 package. The graphical output was produced by ggplot2 package v. 3.3.5 and Pheatmap v. 1.0.12.

### 2.6. Cell proliferation assay

The human colon adenocarninoma Caco-2 cell line was purchased by the Italian Biobank of Veterinary Resources and routinely maintained in Eagle’s minimum essential medium supplemented with sodium pyruvate (1 mM), 1% glutammine and 10% fetal bovine serum. For the cell proliferation assay, cells were seeded at 5E4 cells/mL into 96-well culture plates (100 uL) 24 h before exposure to allow them to adhere to the wells. The cells were then incubated for 24, 48 and 72 h with different concentrations of co-composting elutri-ates in water and DMSO was used as a control vehicle for the extracts at final concentrations never exceeding 1% v/v. Cell proliferation was assessed every 24 h by the MTS assay (CellTiter 96® AQueous One Solution Cell Proliferation, Promega). DMSO-treated cells were used as negative controls and the effects on cell proliferation were expressed as the percentage inhibition compared to the cells of the control group. The related statistical analysis to describe the dose response relationship was performed on R 4.1.2 using Drc package (v. 3.2-0) with the appropriate regression model for each time point. The half maximal effective dose (ED50) and p-value were established using: the Brain-Cousens hormesis models (BC.5) for T0, the four-parameter Weibull functions (W1.4) for T18 and the asymptotic regression model (AR.3) for T28. In each function the lower limit was fixed to “0”.

### 2.7 Micronucleus frequency assay

Seeds of *Vicia faba* L. were germinated at 24± 1 C° for 72 h directly on co-composting sediments or on co-composting sediments elutriates in water. In each experiment, the negative control was achieved using distilled water. After 72 h of germination in the different matrices (control = water; T0 = elutriate from the co-composting sediments at the time of mesocosms set up and the corresponding co-composting sediments; T18 = elutriate from the co-composting sediments after 18 days of incubation and the corresponding co-composting sediments; T28 = elutriate from the co-composting sediments after 28 days of incubation and the corresponding co-composting sediments), the Micronucleus frequency assay (MNC test, number of micronuclei per 1000 nuclei) was adopted for the evaluation of the genotoxicity of the matrices.

## 3. Results

### 3.1 Quantification of the Total Petroleum Hydrocarbons, Humic and Fulvic Acids and Ergosterol content in Mesocosms

The presence of the metabolically active *Lambertella* sp. MUT 5852 was accompanied by a complete depletion of TPH content after 28 days of incubation (Figure 1, panel A). Any decrease in TPH was observed in the co-composting sediments in presence of the autoclaved *Lambertella* sp. MUT 5852 (data not shown). The content of ergosterol in the co-composting sediments showed a continuous decrease with time of incubation (Figure 1, panel B). Humic and fulvic acids did not increase significantly during the co-composting process (Figure1, panels C and D).

**Figure 1.**
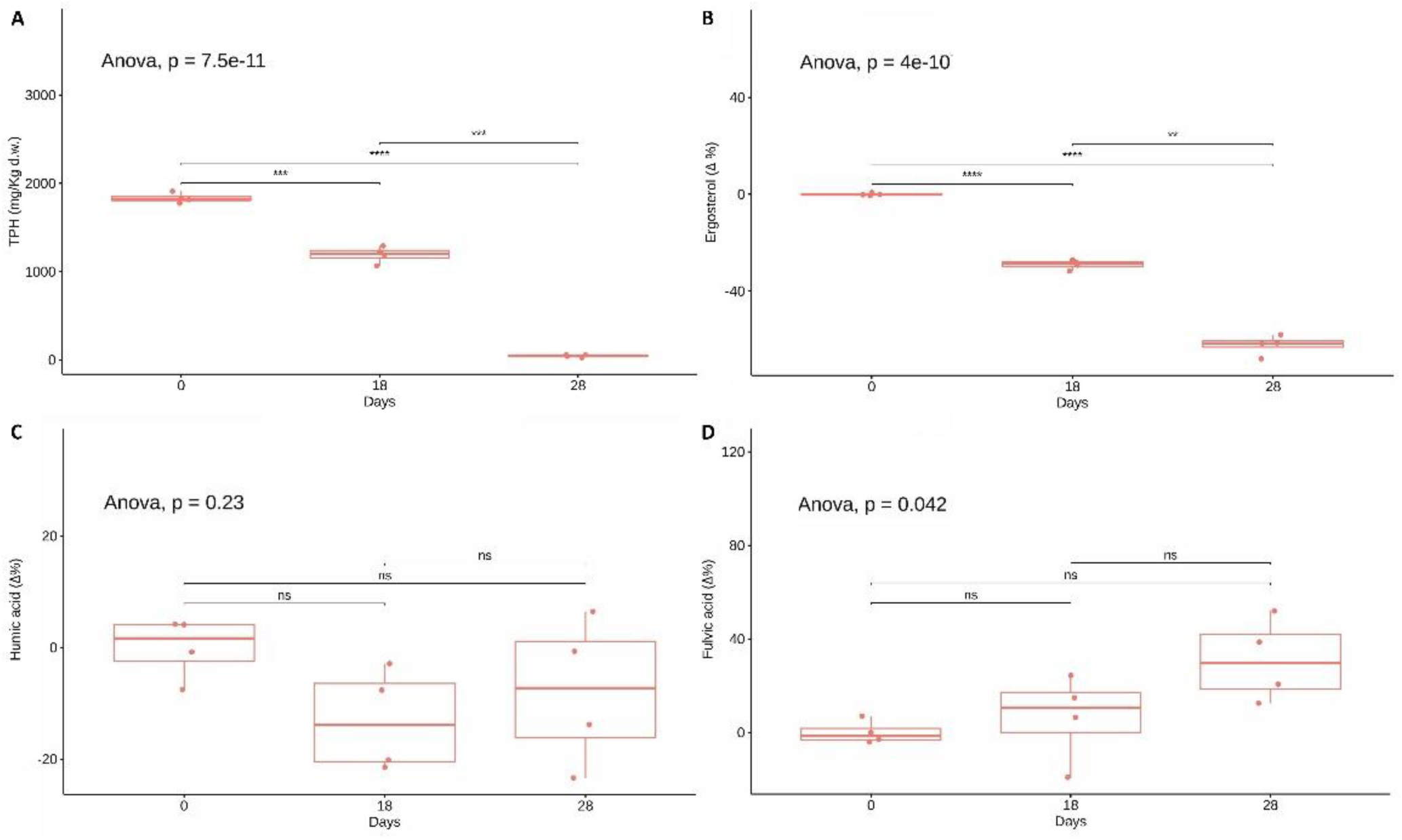
Changes in chemical parameter values during the co-composting process. Val-ues are expressed as part per million (ppm) for TPH and as delta in percentages for the other parameters. Three biological replicates per group are reported. Box and whiskers represent the minimum (Q0), 1st quartile (Q1), median (Q2), 3rd quartile (Q3), and maximum (Q4) of each group. Global statistical significance was evaluated by one-way ANOVA. Horizontal bars show statistical significance of multiple comparisons, calculated by post hoc paired t-tests between group means. Notation for p-value: ns for p > 5 × 10−2; * for p ≤ 5 × 10−2; ** for p ≤ 1 × 10−2; *** for p ≤ 1 × 10−3.

### 3.2 Taxonomic and functional analysis

Results related to the statistically significant changes in relative abundances of bacterial taxa during the process of TPH depletion, calculated by the bias correction in microbiome composition analysis, are reported in Figure 2.

**Figure 2.**
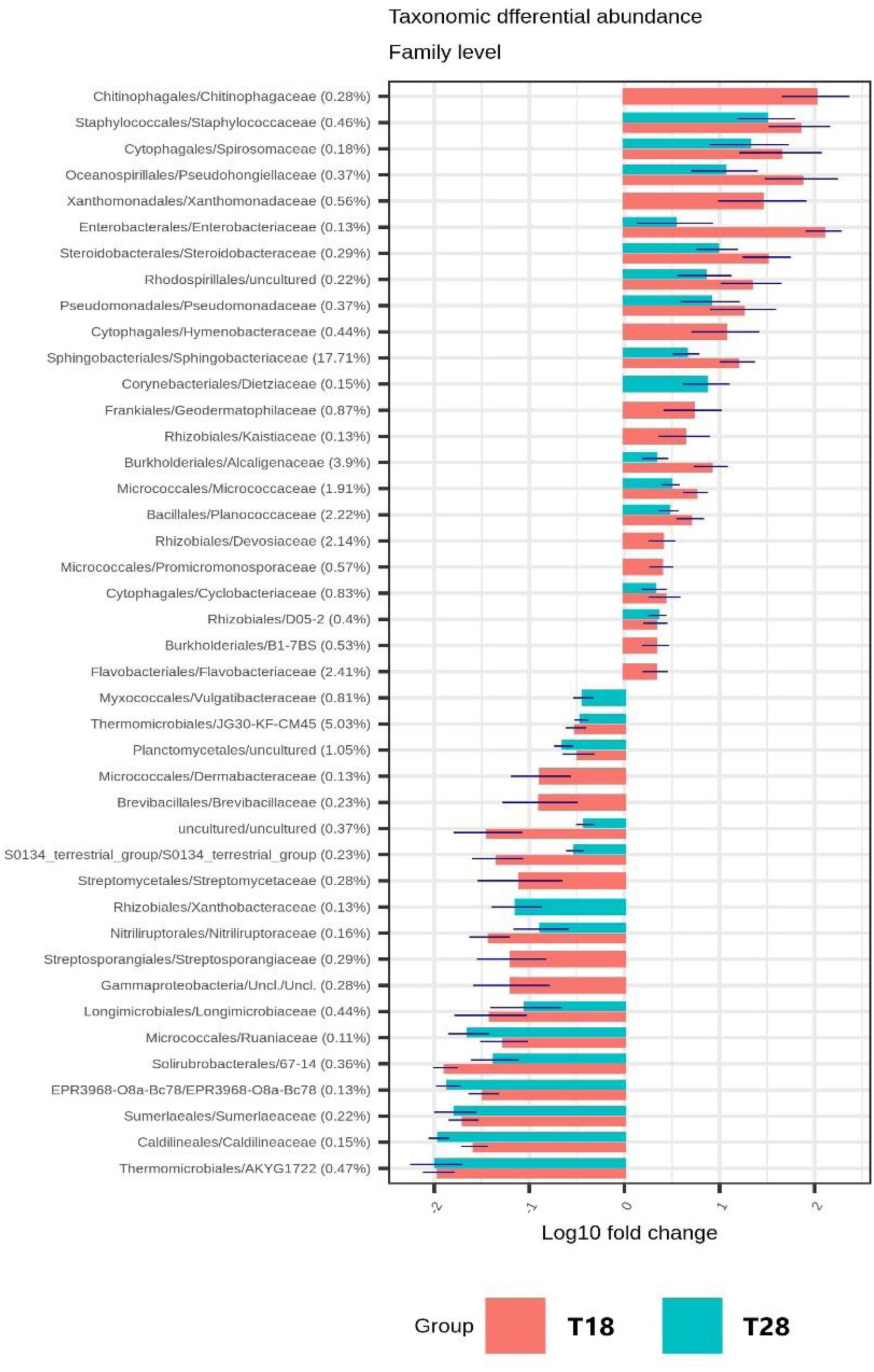
Grouped barplot showing the taxonomic differential abundance at family level between: T18 and T0 in orange, and T28 e T0 in light blue. Statistical results obtained using Analysis of compositional data with bias correction approach (ANCOMBC). Percentage reported near ASV names represent the relative abundance of the sum of the corresponding taxa for ASV against the ASV total sum, without any cutoff.

A decrease in relative abundances of up two order of magnitude was observed at T18 and T28 time of incubation, with reference to the time of set up of the mesocosms (T0). At the same time an increase of the same magnitude was observed for diverse bacterial families. The families with the highest increments in relative percentages of representativeness were the *Flavobacteriaceae* (2.41%) increasing at T18, *Devosiaceae* (2.14%) increasing at T18. A parallel increase at T18 and T28 was observed for the *Planococcaceae* (2.22%), *Micrococcaceae* (1.91%), *Alcaligenaceae* (3.9%), *Sphingobactericeae* (17.71%) families.

To better evaluate the metabolic potential of the bacterial ecology during the process of TPH depletion, the contribution of the different bacterial taxa to the abundance of functional features of interest was analyzed, by evaluating the contribution of specific enzymatic commission (EC) numbers. Results obtained are shown in Figures 3-5.

**Figure 3.**
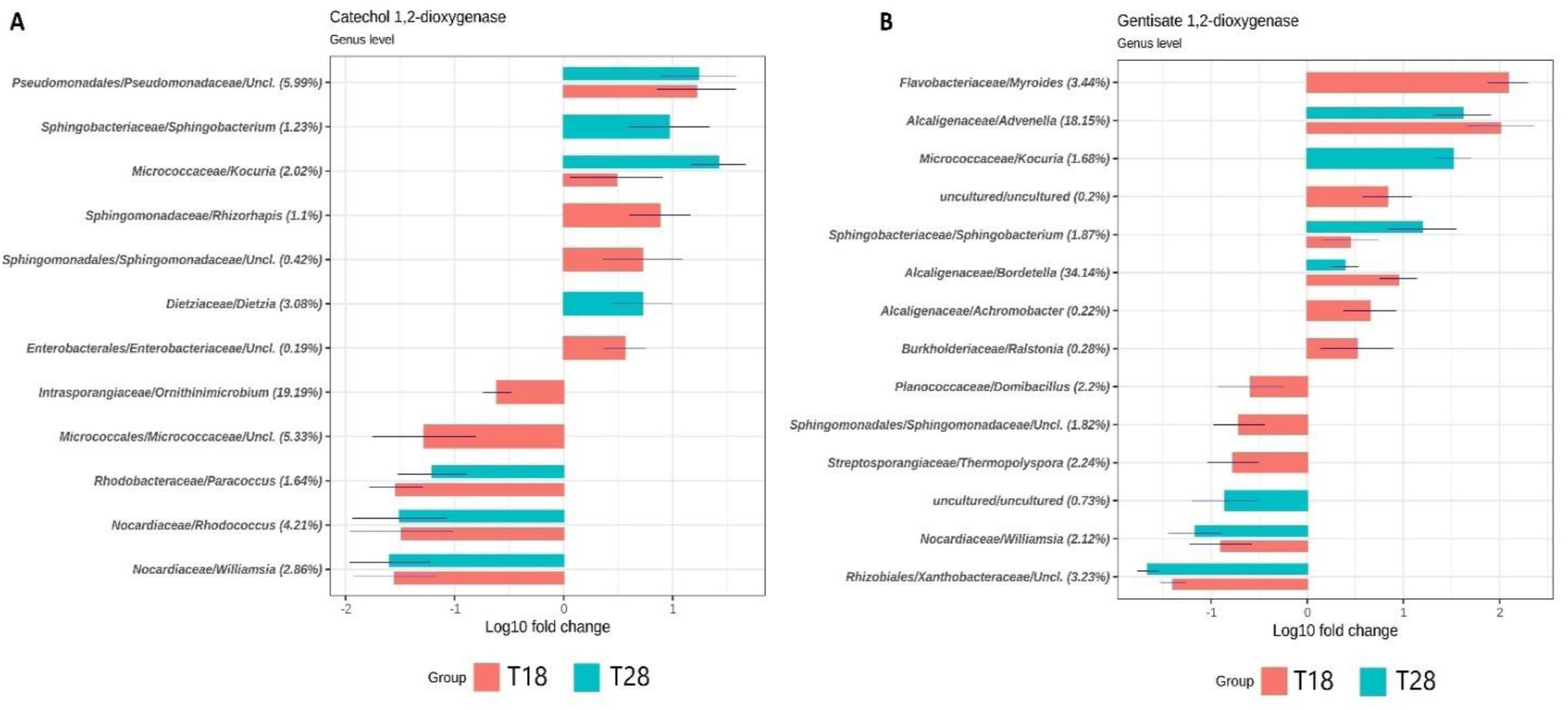
Grouped barplot showing the genus with significantly different abundances between: T18 and T0 in orange, and T28 e T0 in light blue on Catechol 1,2-dioxygenase (Panel A) and Gentisate 1,2-dioxygenase (Panel B). Statistical results obtained using Analysis of compositional data with bias correction approach (ANCOMBC) Percentages reported near ASV names are the relative abundance of the sum of EC count per ASV against that EC total sum, without any cutoff.

Results obtained in relation to the Cathechol 1,2-dioxygenases are reported in Figure 3A. Cathechol 1,2-dioxygenases participate in the lower pathway of aromatic compound degradation via 3- oxoadipate (Pérez-Pantoja et al. 2012), specifically in the ortho-cleavage of the catechol aromatic ring. The highest increases in contribution were recorded at T18 principally in relation to the Pseudomonadaceae family (5.99%) and *Kocuria* sp. (2.02%), both contributing also at T28. *Dietzia* sp. (3.08%) contributed principally at T28.

In relation to Gentisate 1, 2-dioxygenases (Figure 3B), involved in the aromatic ring cleavage of gentisate in m-cresol oxidation (Hopper and Taylor 1975), the increase in the contribution was recorded at T18 with the significant participation of *Bordetella* (34.14%), *Advenella* (18.15%) and *Myroides* (3.44%) sps. *Advenella* sp. increased the contribution also at T28 with Sphin-gobacterium (1.87%) and Kocuria (1.68%) sps..

A further dioxygenase involved in the lower pathway of aromatic compound degradation via 3- oxoadipate, and in the super-pathway of aromatic compound degradation via 2-hydroxypentadienoate (Li et al 2004), is the Catechol 2,3-dioxigenase (Figure 4A). A higher biodiversity with reference to Cathechol 1,2-dioxygenases and Gentisate 1, 2 dioxygenases in bacterial taxa increasing their contribution in this specific enzymatic activity, during TPH depletion, was observed. The evenness in the contribution was high even though *Glutamicibacter* (3.74%), *Sphingobacterium* (1.73%), *Parapedobacter* (21.71%) and *Devosia* (1.37%) sps. showed significantly higher percentages of contribution with reference to the rest of the bacterial taxa involved. The increase in contribution with reference to T0 was similarly distributed among T18 and T28 (Figure 4A). The Protocatechuate 3,4-dioxygenases participates in the lower pathway of a second branch of the super-pathway of aromatic compound degradation via 3-oxoadipate (Shen et al 2005), and in gentisate cleavage via the cleavage of protocatechuate (Figure 4B). The evenness of the increase in contribution on this specific enzymatic activity and the biodiversity of the bacterial taxa involved, were similar to the one observed for Catechol 2,3-dioxygenases even though slightly lower. The highest contribution observed was related to *Advenella* (5.28%), *Bordetella* (1.11%), *Glutamicibacter* (9.4%), *Aquamicrobium* (2.01%) and *Devosia* (5.84%) sps. and the increase in contribution was ob-served both at T18 and T28 (Figure 4B).

**Figure 4.**
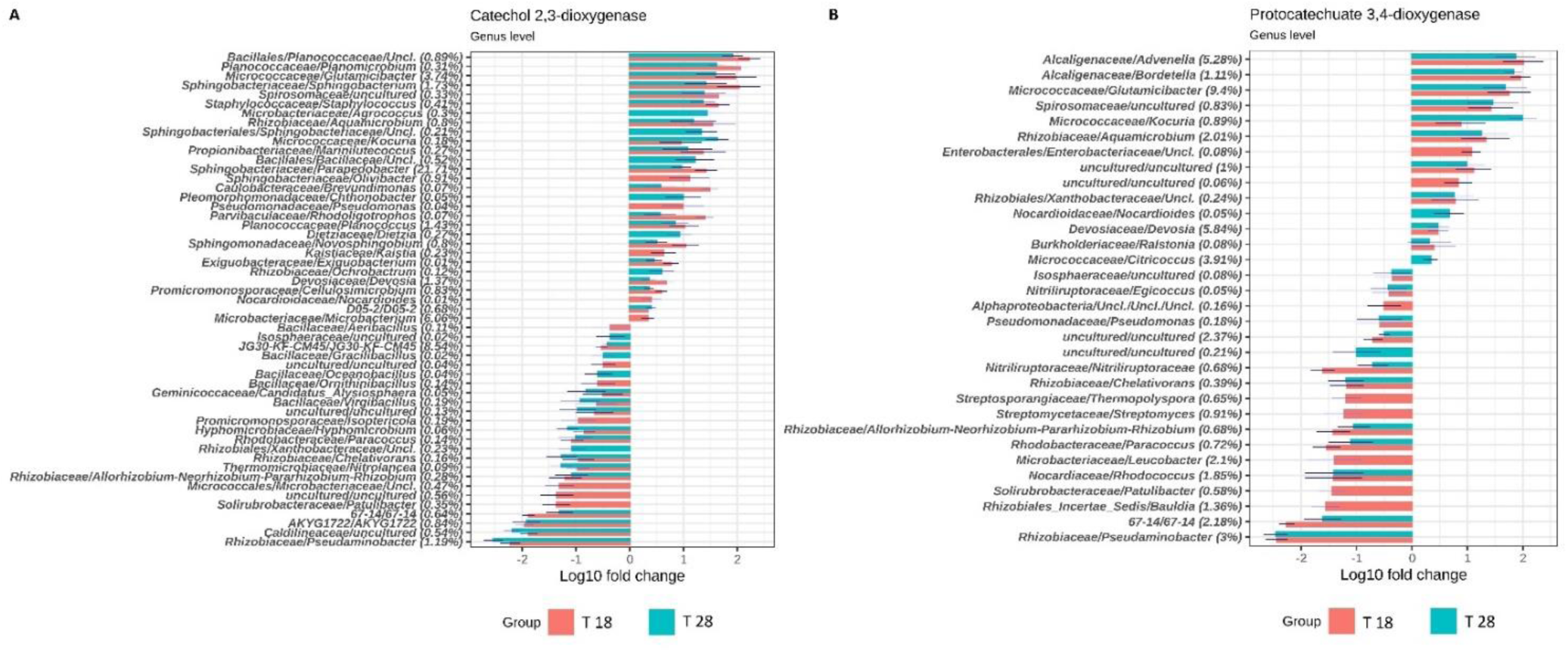
Grouped barplot showing the genus with significantly different abundances between: T18 and T0 in orange, and T28 e T0 in light blue on Catechol 2,3-dioxygenase (Panel A) and Protocatechuate 3,4-dioxygenase (Panel B). Statistical results obtained us-ing Analysis of compositional data with bias correction approach (ANCOMBC) Percent-ages reported near ASV names are the relative abundance of the sum of EC count per ASV against that EC total sum, without any cutoff.

For all the oxidizing activity analyzed, significant decreases in contribution of a plethora of bacterial taxa at T18 and T28, with reference to T0, were observed. It is reasonable to assume that, even though potentially competent for TPH depletion, these bacterial taxa were not contributing to the process and for the same reason they were not described in detail.

In relation to the concept of soil as a matrix offering ecosystemic services, the increase in the contribution of bacterial species capable to promote plant growth occurring in parallel to the process of TPH depletion was also evaluated. In the co-composting sediments, the contribution was principally related to the 1-aminocyclopropane-1-carboxylate (ACC) deaminase activity, which was analysed, since the enzyme inhibit the production of the plant stress hormone, the ethylene, involved in plant stress response to the environment (Penrose et al 2003) (Figure 5). The most relevant contributors were the *Pedobacter* (53.35%) and *Sphingobacterium* (12.55%) sps., both at T18 and T28. The biodiversity of the bacterial genera contributing to the function is relevant. The increase in contribution was equally distributed between T18 and T28, except for *Olivibacter* (2.27%) and *Myroides* sps., which showed a significant increase in their contribution only at T18.

**Figure 5.**
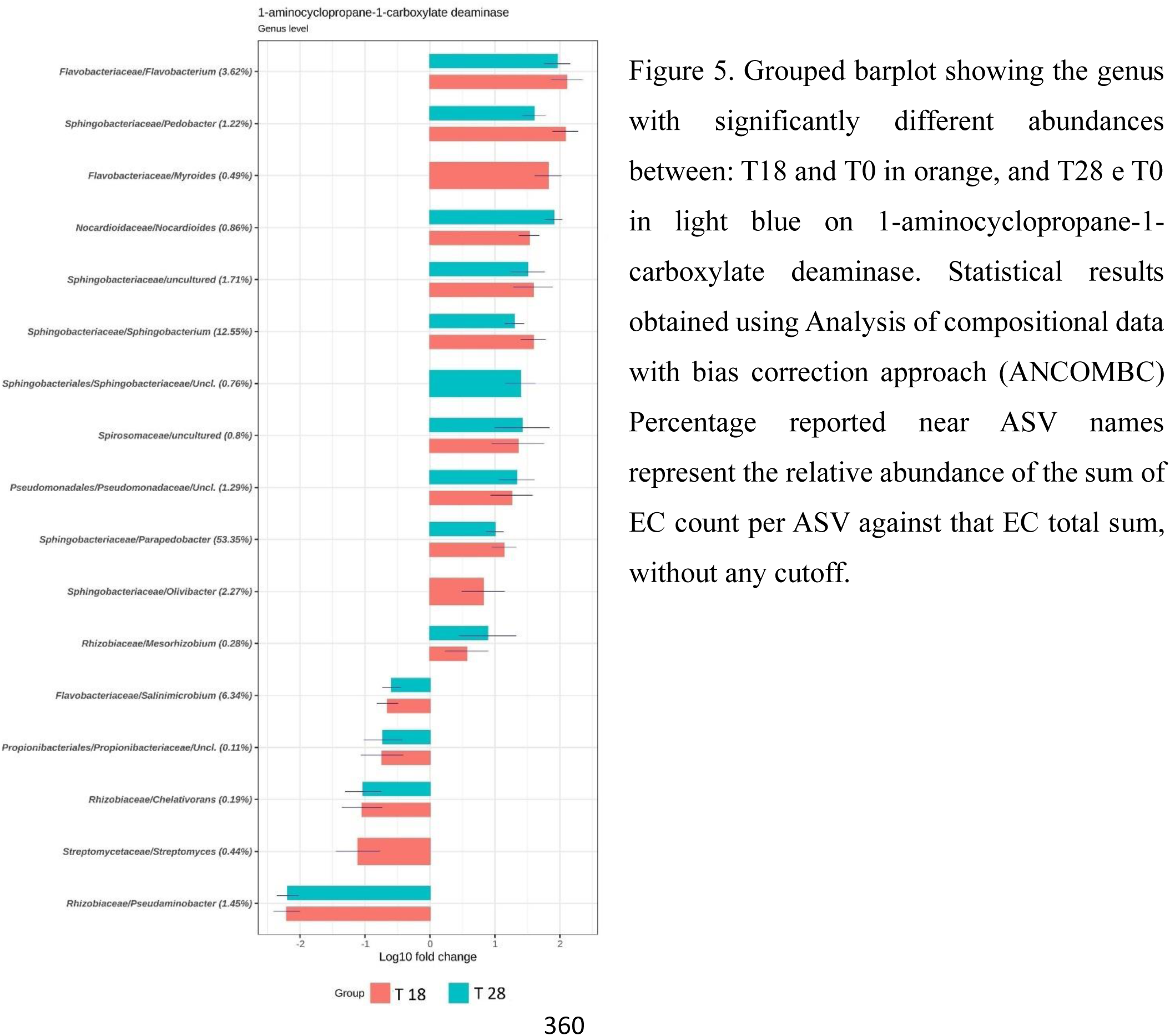
Grouped barplot showing the genus with significantly different abundances between: T18 and T0 in orange, and T28 e T0 in light blue on 1-aminocyclopropane-1-carboxylate deaminase. Statistical results obtained using Analysis of compositional data with bias correction approach (ANCOMBC) Percentage reported near ASV names represent the relative abundance of the sum of EC count per ASV against that EC total sum, without any cutoff.

At the same time, among the metabolic traits related to a resilient soil capable of ecosystemic services, it is relevant the soil microbial capacity to transform the organic matter by humification processes and one the bacterial enzyme involved in the process there is the Dye decolorizing peroxidases (Fig. 6) responsible for the extracellular oxidation of phenolic compounds [17,18], *Micrococcaceae* were principally contributing to the activity, with the specific intervention of the *Glutamicibacter* sp. representing the 15.26% on the total of corresponding EC, in the frame of the total bacterial community contribution to the enzymatic activity. The increase was observed both at T18 and T28. Moreover, within the Micrococcaceae family, *Kocuria* sp. contributed with the 1.44% of the total contributions. Nocardiaceae contributed with the *Gordonia* sp. (2.24%), showing an increment in contribution both at T18 and T28. An increment in contribution was observed at T18 for *Micromonospora* sp. (0.1%).

**Figure 6.**
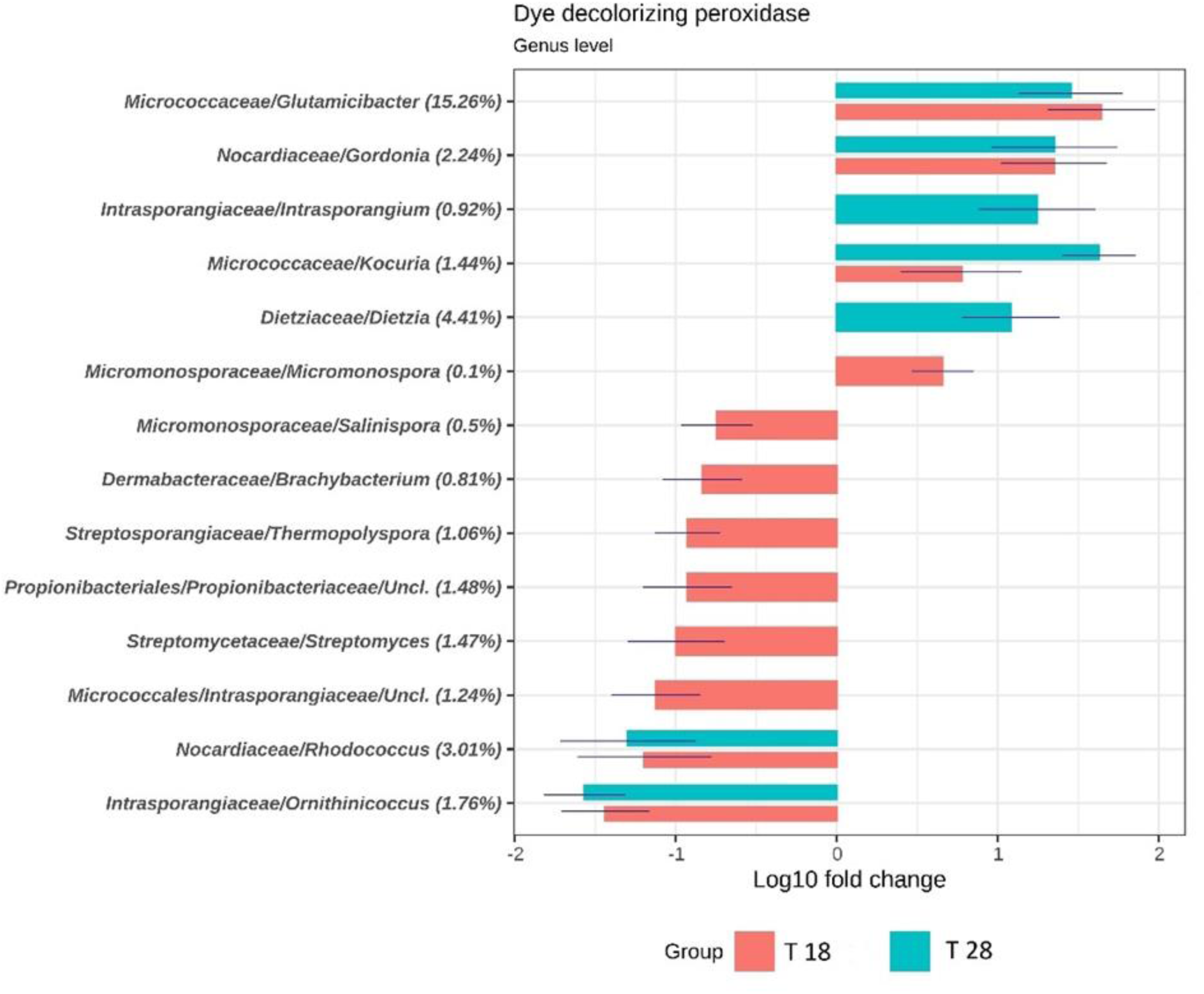
Grouped barplot showing the bacterial genera with significantly different abundances between: T18 and T0 in orange, and T28 e T0 in light blue on Dye decolorizing peroxidase (Panel A) and Alkane 1-monooxygenase (Panel B). Statistical results obtained using Analysis of compositional data with bias correction approach (ANCOMBC) Percentages reported next to the ASV names are the relative abundance of the sum of EC count per ASV against that EC total sum, without any cutoff.

### 3.3 Plant Bioassay

The genotoxicity of the co-composting sediments along the bioremediation process was estimated by analyzing the frequency of micronuclei in interphase of the *Vicia faba* root meristematic cell population, when incubated to germinate in the co-composting sediments and in the corresponding elutriates in water, at the different time of the experimentation. Results obtained (Figure 7) showed a higher genotoxicity of the co-composting sediments with reference to their elutriates in water. The toxicity of the co-composting sediment elutriates (Figure 7, panel A) resulted to continuously decrease with time of incubation, however, it was low even at the beginning of the experimentation. On the other hand, the genotoxicity of the co-composting sediment resulted to be high at T0 (Figure 7, panel B). This latter decreased with time of incubation, even though a residual genotoxicity of the co-composting sediments was observed at the end of the experimentation.

**Figure 7.**
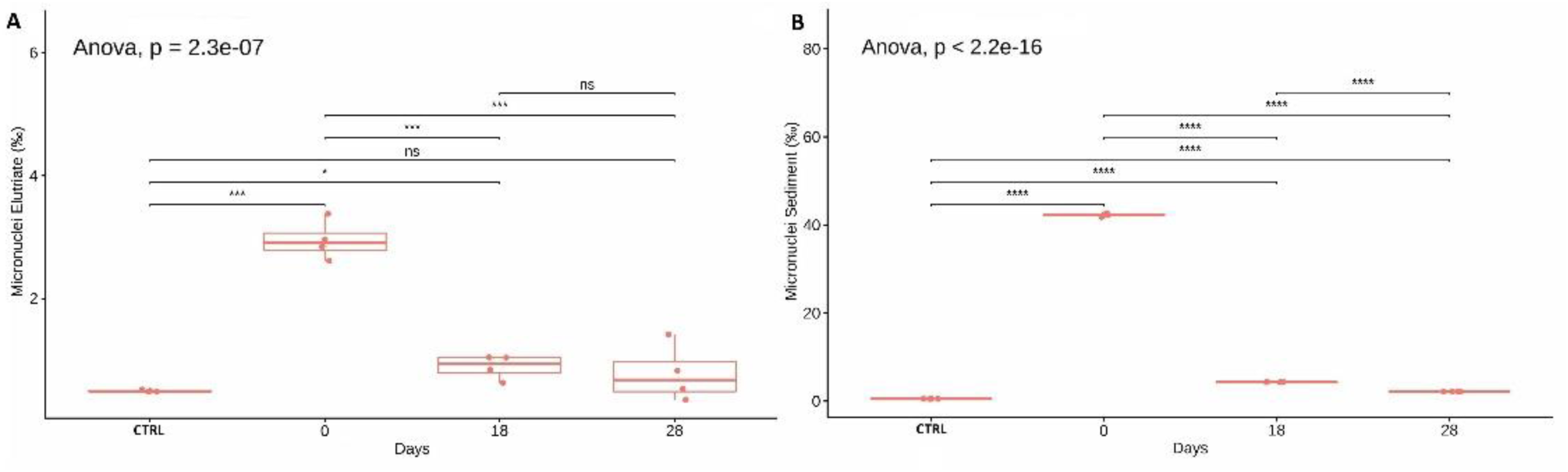
Variation of Micronuclei counts on one thousand cells counted. Reported box-plots represent number of micronuclei variations from the beginning of the experimenta-tion, expressed as cell with micronuclei on one thousand cells of root tip exposed to elu-triates of the co-composting sediments in water (Panel A), to the co-composting sedi-ments (Panel B). The control, CTRL, consists in root tip exposed to tap water. Four biolog-ical replicates per group are reported. Box and whiskers represent the minimum (Q0), 1st quartile (Q1), median (Q2), 3rd quartile (Q3), and maximum (Q4) of each group. Global statistical significance was evaluated by one-way ANOVA. Horizontal bars show statis-tical significance of multiple comparisons, calculated by post hoc paired t-tests between group means. Notation for p-value: ns for p > 5 × 10−2; * for p ≤ 5 × 10−2; ** for p ≤ 1 × 10−2; *** for p ≤ 1 × 10−3.

### 3.4 CaCo-2 cell proliferation assay

The toxicity of the co-composting sediment elutriates in water at the different time of the experimentation was evaluated also on the proliferation of the human intestinal tract CaCo-2 cells (Figure 8), by the estimation of the median effective dose (ED50) associated to the inhibition of cell proliferation. Data are fitted with the proper regression model: the Brain-Cousens hormesis models for T0, the four-parameter Weibull functions for T18 and the asymptotic regression model for T28. In each function the lower limit was fixed to “0”. ED50 and corresponding p-values are reported in each panel of Figure 8. Results obtain showed that the co-composting sediment elutriate showed a significant inhibition of cell proliferation at the highest concentrations. However, the elutriates corresponding to T18 (Figure 8, panel B) showed a certain level of cell proliferation inhibition also at the lowest concentration. In fact, a maximum of 80% of cell proliferation with reference to the inter-nal control was observed. On the other hand, the elutriate at T0 showed an hormetic effect, with the increase in concentration, consisting in a stimulatory effect of cell prolifera-tion, followed by a successive decrease of the corresponding value (Figure 8, panel A). The values of the ED50 at the different time of incubation indicated that the elutriate at T18 showed the highest toxicity (ED50=11.61) with reference to T0 that showed a lower ED50 (20.82). The lowest ED50 was recorded for the elutriate at T28 (121.5), showing a depletion of the toxicity of the elutriates during the co-composting process.

**Figure 8.**
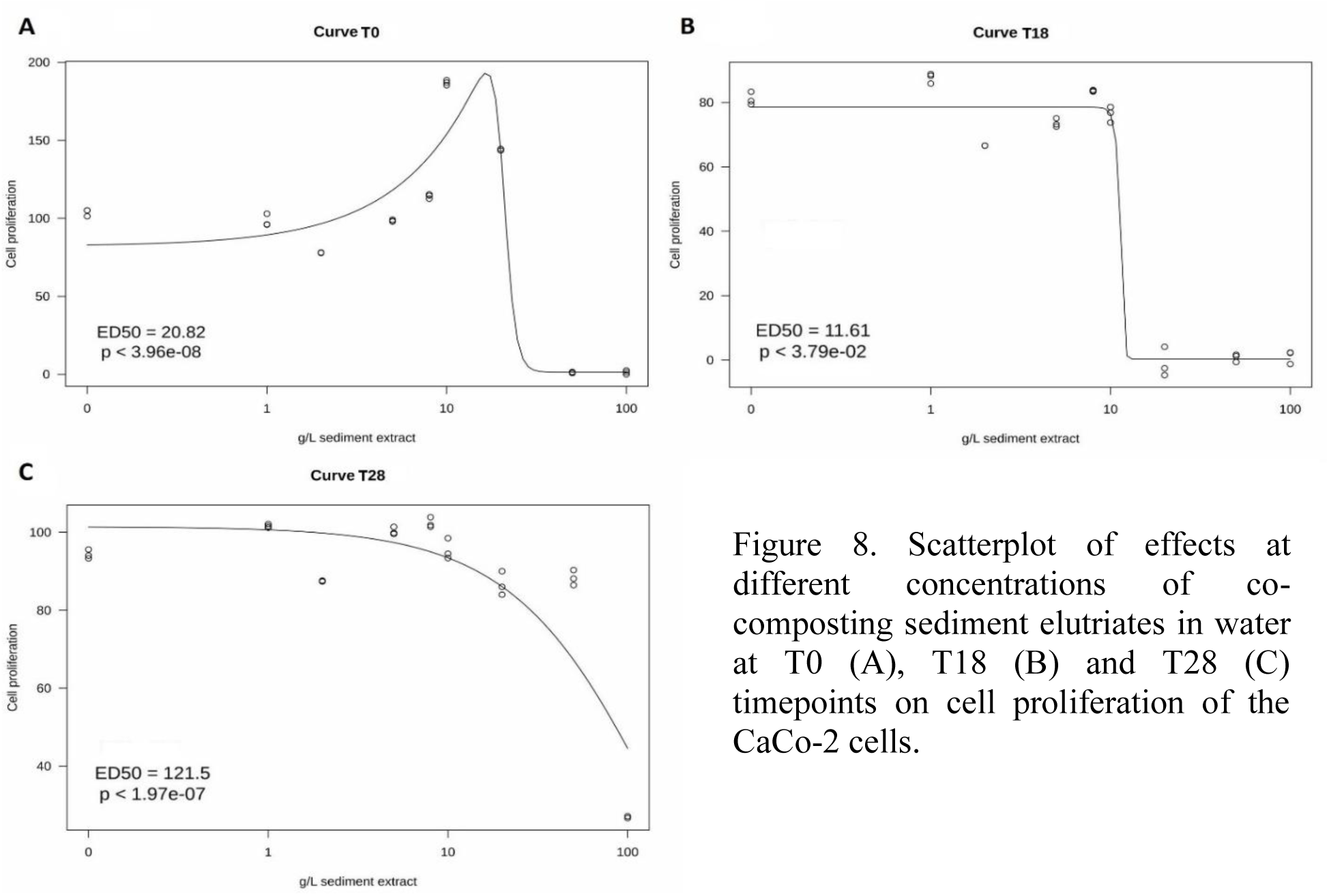
Scatterplot of effects at different concentrations of co-composting sediment elutriates in water at T0 (A), T18 (B) and T28 (C) timepoints on cell proliferation of the CaCo-2 cells.

### 3.5 Canonical correspondence analysis of the process

In CCA biplot (Figure 9), the disposition of datapoints toward the unconstrained varia-bles (TPH, ergosterol content and micronuclei frequency,) whose values showed significant differences with reference to the incubation period, evidenced that the bacterial diversity changed significantly during the co-composting process and that diverse bacterial population were associated to significantly diverse values of unconstrained variables. More precisely the bacterial diversity of T0 and T18 are positively correlated with the higher values of TPH concentration and toxicity. However, the bacterial diversity at T18 is significantly diverse from the one at T0. The differences are evident also for the bacterial diversity of the co-composting sediments at T28, which correlated with the lower values of the unconstrained variables.

**Figure 9.**
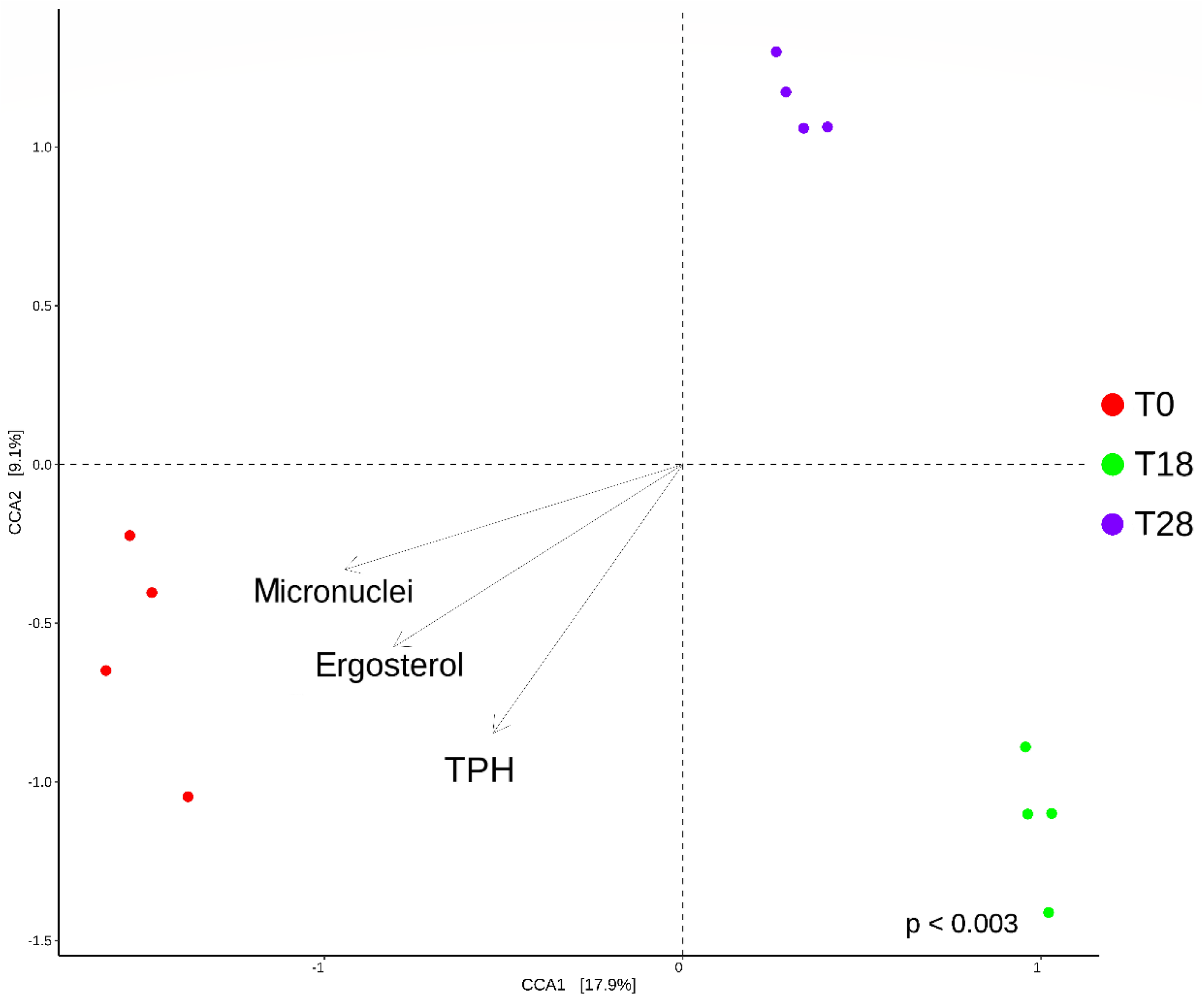
Canonical correspondence analysis (CCA) of the co-composting process. Canonical correspondence analysis (CCA) biplot shows correlation between ASV composition (bacterial ecology) of each sample replicate and the environmental parameters: Micronuclei (referring to the co-composting sediments), TPH, and ergosterol. Colors indicate time category for each datapoint. Black arrows are the eigenvectors representing constraining variables. Reported p-value is calculated by PermANOVA test performed on a full model with 999 permutations.

## 4. Discussion

The co-composting of TPH contaminated dredged sediments with lignocellulosic residues, in combination with the bioaugmentation of the ascomycetes *Lambertella* sp. MUT 5852, already showed encouraging results (Becarelli et al 2021a). However, the success of the approach is related to its transferability on the real scale, and the sustainability in term of cost of the process is mandatory. In this context it is mandatory to reduce the costs of production of the biomass for bioaugmentation. Thus, the decrease of the biomass to be bioaugmented is desirable. Results here obtained showed that the decrease of the density of the inoculum up to the 50% with reference to the previous experimentation is efficient in determining favorable kinetics of depletion of the contamination. Moreover, as observed in (Becarelli et al 2021a), the bioaugmentation of *Lambertella* sp. MUT 5852 is mandatory for the depletion of the contamination, not observed in absence of the fungal inoculum. Despite this assessment, the quantification of ergosterol as marker of the active fungal metabolism (Buiarelli et al 2013; Olsson et al 2003) suggests a decrease of the fungal activity with time of incubation. In the case of bioaugmentation, the decrease in the metabolic activity of the inoculated strain, is a desirable goal at the end of the process of decontamination, to avoid the dominance of the inoculum, over the rest of the microbial community, and the loss of the microbial evenness associated to soil resilience. On the other hand, results here obtained suggest that the Ascomycetes is participating at the depletion of TPH in soils, interfacing with the bacterial community indigenous to the matrices and competent for the transformation of the contaminants. More precisely, in the present experimentation the depletion was the consequence of a synergic activity of the *Lambertella* sp. MUT 5852 and the bacterial community indigenous to the sediments, as suggested by the increments in relative abundances of specific bacterial taxa specialized in the depletion of the contamination (specialist species). Their intervention in the degradation of TPH was assessed by the harboring of enzymatic activities such as the catechol 1,2-dioxygenases, gentisate 1,3-dioxygenase, catechol 2,3-dioxynaes and protocatechuate 3,4-dioxygenases. In fact, these enzymes are all in-volved in the lower pathways of the oxidation process of aromatic chemical structures or the ring cleavage of aromatic acid derivatives (Ferraro et al 2005). In this context it should be mentioned that TPH are characterized the unresolved complex mixture (UCM), the complex chemical fraction responsible for the toxicity of the contamination (Scarlett et al 2007; Thomas et al 1995), which includes thousands of saturated and unsaturated compounds, comprising polyaromatic chemical structures recalcitrant to biodegradation (Booth et al 2007; 2008). It is a consolidated awareness that these compounds cannot be identified by established target analysis (Samanipoura et al 2015). The same can be assessed for the intermediates of degradation of high molecular weight polyaromatic structures. These molecules are partially oxidized aromatic structures with decreasing molecular weights, because of the progressive cleaving of composing aromatic rings. The dioxygenases here contributing to the depletion of TPH might be responsible for these cleavages, funneling the transformation of the recalcitrant to biodegradation structures and the net depletion of the TPH. The microbial genera here observed as increasing in their contribution during the co-composting process, have been already described as harbouring the enzymes involved in the degradation of the polycyclic aromatic hydrocarbons, reported also as composing the UCM (Booth et al 2007; 2008): e.g.: *Kocuria* (Li et al 2016), *Dietzia* (Hidalgo et al 2020), *Bordetella* (Eriksson et al 2003), *Advenella* (Wang et al 2014), *Myroides* (Maneerat et al 2006), *Glutamicibacter* and *Sphingobacterium* (Lu et al 2019), *Parapedobacter* (Wu et al 2020), *Devosia* (Song et al 2021), *Aquamicrobium* (Andreaoni et al 2004) sps.. More precisely, all the taxa were described for the oxidation of the intermediates of degradation of polyaromatic structures. Thus, in relation to the interplay with *Lambertella* sp. MUT 5852 during the co-composting processes, the described bacterial taxa represent the specialist species directly involved in the degradation of the contamination. The changes in their relative abundances, that determine the changes in the bacterial biodiversity during the process of TPH degradation, resulted to be strongly correlated to the kinetics of TPH depletion as shown by the canonical correspondence analysis. More precisely, they are responsible for the oxidation of intermediates of degradation of the primary pollutants, eventually partially oxidised by the extracellular oxidising enzymatic battery of the bioaugmented ascomycetes (Zavarzina et al 2010) and/or of other oxidizers activated by the co-composting process (Tran et al 2021).

In addition to the decontamination, the recovery of contaminated dredged sediments as recycled materials, should be associated to their safety in toxicological terms at the end of the treating process. In this context, the bacterial enzymatic activity analyzed results to be pivotal to the depletion of eventual toxic intermediates of degradation of the primary pollutants, participating both to the process of decontamination and detoxification of the matrix in treatment. In fact, a strong correlation was observed also between the bacterial biodiversity changes during the process of decontamination and the decrease in genotoxicity of the co-composting sediments. Thus, the observed changes in bacterial diversity are associated to both the decontamination and detoxification of the matrices. More in details, the increment in dominant bacterial taxa at T18 suggests the latter as mandatory to determine the kinetic of TPH depletion observed, since they characterize the co-composting sediments in the process phase associated to the maximum slope of the TPH depletion kinetic. The lasting of the contribution of the same bacterial taxa at the end of the experimentation, is related to a residual presence of intermediates of degradation, which are responsible for the residual genotoxicity measured in the co-composting sediments at the end of the experimentation and/or for the toxicity assayed by the CaCo-2 cell proliferation of the co-composting sediments elutriates. On the other hand, the observed decrease in contribution of bacterial taxa harboring the enzymatic activity of interest, all over the experimentation, is related to the evidence that the dioxygenases activities, responsible for TPH depletion, are associated to the specificity for the substrates of the enzymes involved (Ferrero et al 2005). In this context it is reasonable to assess that the chemical structures of the intermediates of degradation of the primary pollutants and their concentrations, are responsible for the increment in specific bacterial populations, harboring competent enzymes, with the associated depletion of bacterial populations not competent for the same chemical structures.

In relation to the DyP, the enzyme has been described for the extracellular oxidation of chemically complex phenolic polymers by the a-specificity for the substrates of the enzyme (Chen et al 2016; Chen et al 2015). Due to the characterizing high redox potential, the enzyme might be responsible for the oxidation of the primary pollutants and their intermediates of degradation. However, in similar process conditions, the enzyme has been described as associated to the saprophytic metabolisms of bacterial generalist species, interacting the fungal one, for the increment in bioavailability of the contamination (Becarelli et al 2021b). Due to the a-specificity of the catalytic activity of the enzyme, a process of speciation of the harbouring bacterial community, induced by the chemical structures of contaminants is not consistent. The high redox potential extracellular oxidases, such as the DyP, are responsible for polycon-densation reactions, involved in the humification of the organic matter (Wang et al 2008). Even though a participation of the DyP to the oxidation of the primary pollutants and their intermediates of degradation cannot be excluded, both previous assumptions suggest that the DyP activity might be not directly or not solely involved in the depletion of the contamination, but also in the already described interplay of saprophytic microorganisms in the rearrangement of the organic matter in the substrate. In this context, it should be mentioned the possible involvement of the enzymatic activity in the carbon cycle in soilw, by the capability to mobilise carbon sources through the partial oxidation of humified organic matter at high molecular weight and the release of eventually more bioavailable organic mole-cules, for the nourishment of soil colonisers, participating to the conservation of the soil biodiversity. The metabolic function is of outmost interest in the context of the recovery of a technosol as a resilient matrix. Actually, compost find application in a plethora of noxious conditions, related to both organic and inorganic contamination (Huang et al 2016), since rich in humic matter and microorganisms that interact with the latter, by the enzymatic machinery responsible for the carbon cycle in soil. The recovery of the activity among the bacterial taxa whose relative abundance increased during the process of TPH depletion, confirmed the effectiveness of the process on different aspects, such as the decontamination and detoxification of the matrix and the recovery of metabolic traits associated to the capacity of the latter to offer ecosystemic services.

Noteworthy, the results here obtained showed also the increment in contribution of bacterial taxa harboring the ACC-deaminase, with positive effect on the capability of the colonizing microbial community to promote processes of vegetation, important for the exploitation of decontaminated dredged co-composting sediments as techosols. In fact, the dominant bacterial taxa in functional contribution, *Pedobacter* (Leontidou et al 2020) and *Sphingobacterium* (Chen et al 2022) sps. have been already described as harboring the ACC-deaminase, playing a key role in alleviating plant-stress in diverse environmental conditions.

As a whole, the here described co-composting process induced a significant increment in diverse bacterial taxa offering important metabolic traits for a matrix that should be re-covered to produce technosols. Noteworthy is the co-occurrence of metabolic traits of interest in the same genus such as the case of *Kocuria* sp., which combined all the oxidative functions here analyzed. In addition to the hydrocarburoclastic capacity, the genus is described as highly adaptable to changing and stressful environment (Gholami et al 2015; An et al 2022). At the same time the genus *Sphingobacterium*, in addition to the ACC-deaminase activity, contributed to all the oxidative activities except for the DyP and the Protocatechuate 3,4-dioxygenase. *Sphingobacterium* sp. has been described as playing a key role in affecting the rate of organic matter mineralization in soil, even in stress condition (Rodríguez-Berbel et al 2021; Zhang et al 2021). The same bacterial taxa also determine a decrease in the toxicity of the co-composting sediments. The adoption of two different system for assaying toxicity, with diverse endpoints, evidenced the importance of a multivariate analysis in toxicology, since the micronuclei test resulted to be not sufficiently accurate in assaying the toxicity of the co-composting sediments elutriates, during the treatment. In fact, the sustainability of the technology should be consistent not only with the final product, but also with the process leading to this latter. Toxicological assays provide an instrument to mitigate the noxious effect the process causes to the environment. In this context, the approach here adopted evidenced the necessity to control the toxicity of both the matrix and of the associated elutriates in water, since these latter are eventually responsible for the transfer of toxicants to waterbodies. Differently to other co-composting processes of contaminated matrices with lignocellulosic matrices, an increment in fulvic and humic acids was not observed in the time interval of the process. This latter might be too short to observe the complex process of the humification of the organic fraction that anyway is desirable, since responsible for the soil resilience to environmental disturbance, comprising the presence of contaminants, which are entrapped by the hydrophobic moieties of the organic matter, with a consequent de-crease in bioavailability and toxicity.

## 5. Conclusions

The co-composting process here optimized, with reference to a previous experimentation, was analyzed combining the chemical assessment of the process of TPH depletion to the evaluation of the sustainability of the process in toxicological terms. The bacterial functional contribution to the process is consistent with a multivariate contribution to both the depletion and the detoxification of the matrix and the increase in metabolic traits of interest for both the transformation of the organic matter in soil and the promotion of plant growth even in stress condition. All these functional features indicate the mycobased co-composting process as efficient for the recovery of the treated dredged sediment to produce technosols. At the same time the synergism between *Lambertella* sp. MUT 5852 and the bacterial ecology of the matrix in the process of TPH depletion, was evident. Two specific bacterial strains, the *Kocuria* and *Sphingobacterium* sps., were recorded as extremely peculiar, since harboring oxidative enzymes, possibly involved in both the de-contamination and detoxification of the soil, and in the transformation of the organic matter in the matrix in treatment. In addition, *Kocuria* sp. resulted to be potentially involved also in the synergism between soil bacteria and plants. Results obtained suggest the opportunity to design dedicated culturomic approaches for the obtainment of bacterial microbiota to implement the described mycoaugmented co-composting process to improve the effectiveness of the process both in the direction of decontamination, detoxification and recover of the dredged sediment as a technosol.

## Author Contributions

S.B., G.S, D.B., M.R.C. G.D.S.: Conceptualization, investigation, validation, data curation, writing original draft; G.B.: Conceptualization, investigation, Data curation, formal analysis; methodology, software; S.D.G.: Conceptualization, validation, data curation, writing, reviewing, and editing, supervision, project administration.

All authors have read and agreed to the published version of the manuscript.

